# Minimal polymerase-containing precursor required for Chikungunya virus RNA synthesis

**DOI:** 10.1101/2025.10.06.680709

**Authors:** David Aponte-Diaz, Abha Jain, Jayden M. Harris, Jamie J. Arnold, Craig E. Cameron

## Abstract

Alphaviruses pose a growing global health threat, with Chikungunya virus (CHIKV) epidemics ongoing and no safe or effective vaccines or antivirals currently available. The CHIKV nonstructural proteins nsP2 and nsP4 encode essential enzymatic activities that represent key targets for antiviral development, yet the biochemical basis of nsP4 RNA-dependent RNA polymerase (RdRp) activity remains poorly understood. Here, we identify a minimal, functional precursor form of nsP4 derived from the nsP3–nsP4 polyprotein (P34) that is active in a cell-based RNA replicon system. Using synthetic, capped mRNAs, we show that cleavage of P34 by the nsP2 protease is required for robust reporter expression, and that a truncated form retaining only the C-terminal 50 residues of nsP3 (CT50-P34) supports near–wild-type replication. Unexpectedly, ubiquitin–nsP4 fusions failed to substitute for P34, likely reflecting the transient expression supported by our RNA-based system. We propose that precursor forms of nsP4 interact with the nsP1 dodecamer at the site of genome replication, where cleavage activates the RdRp and localization within the nsP1 dodecamer maintains nsP4 in its active conformation. Dissociation from the nsP1 dodecamer triggers a conformational switch to an inactive state. Together, these findings establish a tractable framework for interrogation of the assembly, activation, and regulation of the alphavirus polymerase.

## INTRODUCTION

Alphaviruses use a polyprotein strategy to express their genome-encoded non-structural proteins [1–3] . The most abundant precursor is termed P123, as it is a fusion of the non-structural protein 1 (nsP1), nsP2, and nsP3 proteins[1] . Readthrough of the opal stop codon present at the end of the nsP3-coding sequence produces P1234 [4] . nsP4 is the viral RNA-dependent RNA polymerase (RdRp) [5] . Among the earliest proteolytic processing pathways proposed for alphaviruses originated from studies by Hardy and Strauss, using Sindbis virus (SINV) as a model (**Fig. 1A**) [6] . P123 is first cleaved to produce P12 and nsP3, followed by cleavage of P12 to nsP1 and nsP2. In contrast, P1234 is first cleaved to produce P12 and P34. P12 is cleaved to nsP1 and nsP2. However, P34 accumulates and exhibits a half-life of approximately 60 minutes [6–8] . Cleavage of P34 to nsP3 and nsP4 occurs over time, but nsP4 does not accumulate to high levels in cells [6,9], presumably because of the instability caused by its Tyr amino terminus and consequential degradation by the proteasome [9–11] . Later studies revealed an alternative pathway for processing of P1234 to P123 and nsP4 [9,12] . This processing pathway only occurred early during infection and may have been restricted to early times post-infection because precursor forms of the nsP2-encoded protease were required to cleave nsP4 from P1234 [9,12] .

**Figure 1.**
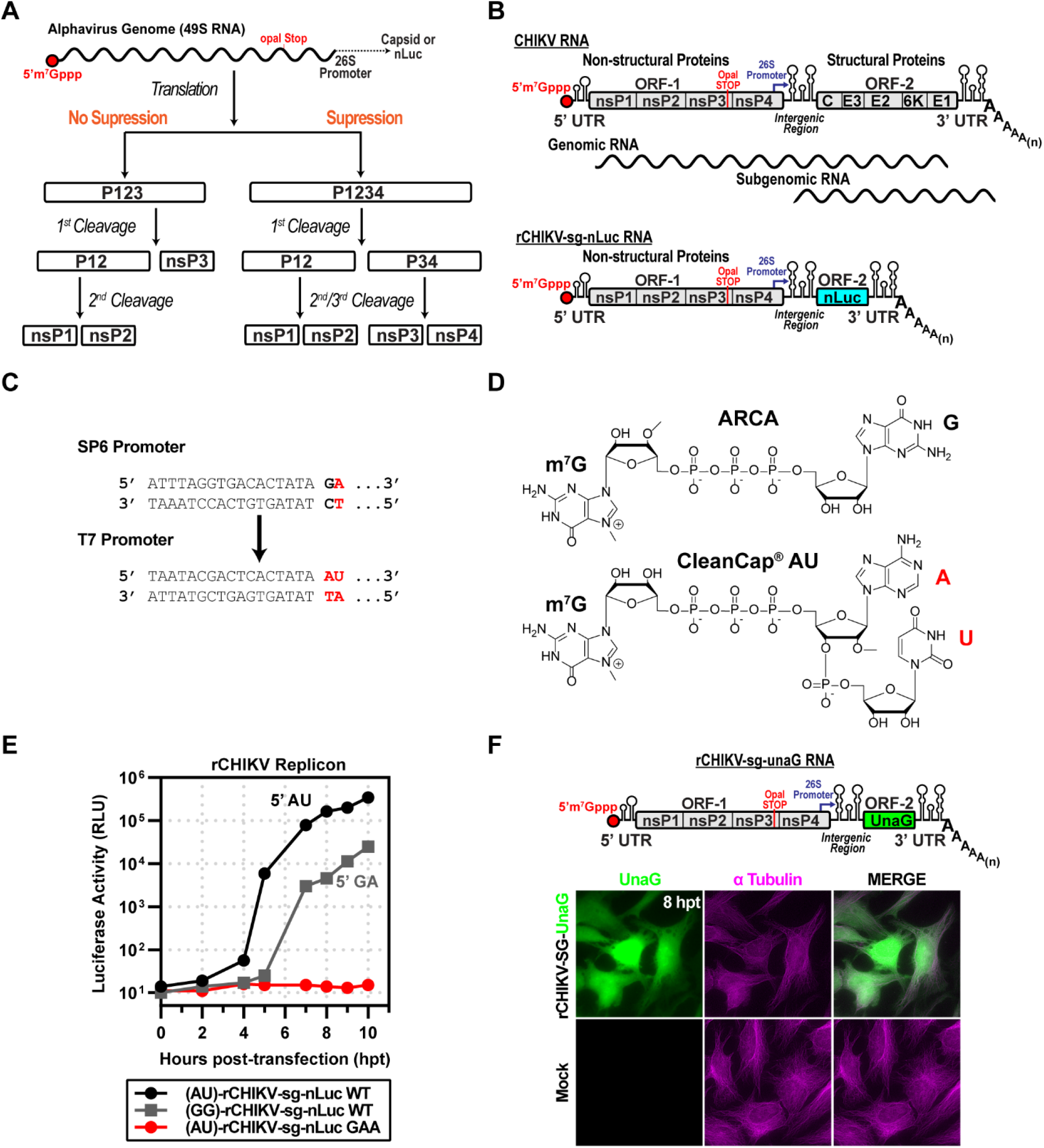
CHIKV replicon used in this study. **(A)** Schematic of proposed alphavirus polyprotein processing pathways. Following entry and uncoating, the genomic RNA (gRNA) undergoes cap-dependent translation through both suppression and non-suppression mechanisms. Without suppression, translation terminates at the leaky opal stop codon at the end of nsP3, producing the P123 polyprotein, which undergoes cis-cleavage, yielding P12 and nsP3. P12 is further processed into nsP1 and nsP2. In the suppression pathway, opal read-through produces P1234, which is cleaved in stages to yield P12 and P34, and subsequently cleaved into nsP1/nsP2 and nsP3/nsP4. **(B)** Organization of the CHIKV genome and replicon (rCHIKV-sg-nLuc). The positive-strand RNA genome is capped at the 5′ end and polyadenylated at the 3′ end. ORF-1 encodes nsP1–4, followed by a 26S subgenomic promoter and ORF-2 encoding structural proteins (C, E3, E2, 6K, E1). The replicon replaces structural proteins with nanoluciferase (nLuc), expressed from the subgenomic RNA. **(C)** Promoter and cap analog comparison: SP6 promoter with ARCA (GA) vs. T7 promoter optimized for CleanCap (AU). **(D)** Chemical structures of ARCA and CleanCap AU used in in vitro transcription of replicons. **(E)** Comparison of capped rCHIKV-sg-nLuc RNAs. AU-capped RNAs yielded stronger luciferase signals relative to GA-capped transcripts. The GAA mutant of nsP4 (polymerase inactive) produced a background signal only. **(F)** Fluorescent reporter version (rCHIKV-sg-UnaG), UnaG placed in ORF-2. Transfected HeLa cells displayed UnaG fluorescence (green), with tubulin immunostaining (magenta) marking cell boundaries. Mock controls lacked UnaG signal.

The observations made by the Strauss laboratory were consistent with the P34 precursor serving as the donor of the nsP4 RdRp required for genome replication and subgenomic RNA synthesis during the exponential phase of the lifecycle [1] . Additional support for the use of the P34 precursor as the donor of the RdRp came from the studies of Lemm and Rice using SINV [7,13] . These studies used vaccinia virus to express precursor and processed forms of SINV non-structural proteins in various combinations. Co-expression of P123 and P34 was both necessary and sufficient to support genome replication and subgenomic RNA synthesis [7,13] . A later study by Lemm and Rice demonstrated that attachment of the carboxy-terminal 22 amino acid residues of nsP3 (referred to herein as CT22-P34) was sufficient to create a form of P34 that appeared to complement P123 as efficiently as the full-length precursor [7] .

Subsequent investigation of CT22-P34 as the donor of the nsP4 RdRp led to the hypothesis that nsP4 had to be expressed in a form that would yield an authentic Tyr amino terminus [9] . To test this possibility, the Rice laboratory fused ubiquitin (Ub) to the amino terminus of nsP4. The resulting Ub-nsP4 fusion paired with P123 to support genome replication and subgenomic RNA synthesis, as well as P34 precursors [12,13] .

The capacity for Ub-nsP4 to substitute for P34 essentially led to the demise of P34. There has been minimal mention of P34, even in the description of the P1234 precursor processing pathway, since the ability of Ub-nsP4 to substitute for P34 was revealed [13] . P34 has been observed in other alphaviruses but has not been studied extensively [14–16] . Indeed, most studies that have evaluated the ability of an RdRp from one species of alphavirus to substitute for that from another species have delivered nsP4 as the Ub-nsP4 fusion [10,11,17] .

We recently reported the purification of a CT50-P34 derivative from O’nyong’nyong virus (ONNV), a close relative of Chikungunya virus (CHIKV), and characterization by small-angle X-ray scattering [18] . This derivative formed a tetramer and concealed the nsP3-nsP4 cleavage site, explaining the previously reported slow cleavage of SINV P34 [6,8,18] .

This study describes the development of RNA-based replicons for CHIKV, which will permit biological analysis of the encoded non-structural proteins. One replicon requires the addition of an mRNA expressing an nsP4 RdRp donor. Unlike the vaccinia- and polII/polI-driven systems reported by others in which the Ub-nsP4 fusion was an efficacious donor of the RdRp, the Ub-nsP4 fusion was a poor RdRp donor in the RNA-based replicon system. Full-length P34, CT150-P34, and CT50-P34 were able to serve as the RdRp donor. We discuss potential reasons for the discrepancy and the implications of our findings on our understanding of alphavirus biology derived from the use of more permissive replicons.

## RESULTS

### CHIKV replicon used in this study

The alphavirus genome encodes two open reading frames: ORF-1 and ORF-2 (**Fig. 1B**). ORF-1 encodes the non-structural proteins responsible for genome replication. ORF-2 encodes the structural proteins responsible for virion morphogenesis. ORF-1 is translated in a cap-dependent manner using full-length genomes, while ORF-2 is translated in a cap-dependent manner from a subgenomic RNA. It is well established that ORF-2 can be eliminated without consequence to the kinetics of genomic or subgenomic RNA synthesis [19] . We substituted ORF2 with nanoluciferase (nLuc)-coding sequence between the genetically encoded start and stop codons (**Fig. 1B**) [20] .

The cDNA used first encoded a bacteriophage SP6 promoter (**Fig. 1C**). Transcription initiation by this RNA polymerase requires that the first nucleotide incorporated is guanosine monophosphate (GMP) [21] . However, the first nucleotide of alphavirus genomes is adenosine monophosphate (AMP) [22] . Therefore, use of SP6 RNA polymerase (SP6) produces an RNA without an authentic 5’-end [21] . We also created a cDNA that utilized a bacteriophage T7 promoter, allowing T7 RNA polymerase (T7) to produce an authentic 5’-end (AU, **Fig. 1C**).

To ensure the addition of an m^7^G cap to the 5’-end, we used cap analogs to initiate transcription [23] . For SP6, we used ARCA; for T7, we used CleanCap® AU (**Fig. 1D**) [24] . We transfected ARCA-capped (5’-GA) or CleanCap® AU-capped (5’-AU) RNA into HeLa cells and measured luciferase activity as a function of time post-transfection. The RNA with an authentic cap replicated faster than one lacking the authentic cap (**Fig. 1E**). Replication of the RNA with the authentic cap required the encoded polymerase, because converting the catalytic site GDD to GAA eliminated RNA synthesis (GAA in **Fig. 1E**).

We also constructed a subgenomic replicon encoding the fluorescent green UnaG reporter [25] (**Fig. 1F**). Transfection of the CleanCap® AU-capped UnaG reporter RNA into HeLa cells led to expression of UnaG (**Fig. 1F**). Colocalization with tubulin was consistent with accumulation of UnaG in the cytoplasm (**Fig. 1F**).

### Strategy to express proteins in cells replicating subgenomic RNA replicons

The goal of this study was to determine the form(s) of CHIKV nsP4 or P34 used for RNA synthesis in cells. We constructed a cDNA that would direct the expression of bicistronic (BiCis) mRNA using T7 and CleanCap® AG. The 5’- and 3’-untranslated regions were taken from human β-globin mRNA, which should confer stability to the RNA (panel i of **Fig. 2A**) [26] . We designed the expression cassette to permit the first cistron (ORF-A) to be changed easily and the second cistron to encode enhanced green fluorescent protein (eGFP in panel i of **Fig. 2A**) [27] . The cap directs translation of ORF-A. Translation of eGFP is driven by encephalomyocarditis virus internal ribosome entry site (EMCV IRES in panel i of **Fig. 2A**) [28] . To optimize transfection conditions, we introduced the coding sequence for the photostable red fluorescent protein, TagRFP-T, into the first cistron (panel ii of **Fig. 2A**) [29] . Transfection of the bicistronic mRNA encoding both fluorescent proteins produced strong signals for both proteins (panel iii of **Fig. 2A**).

**Figure 2.**
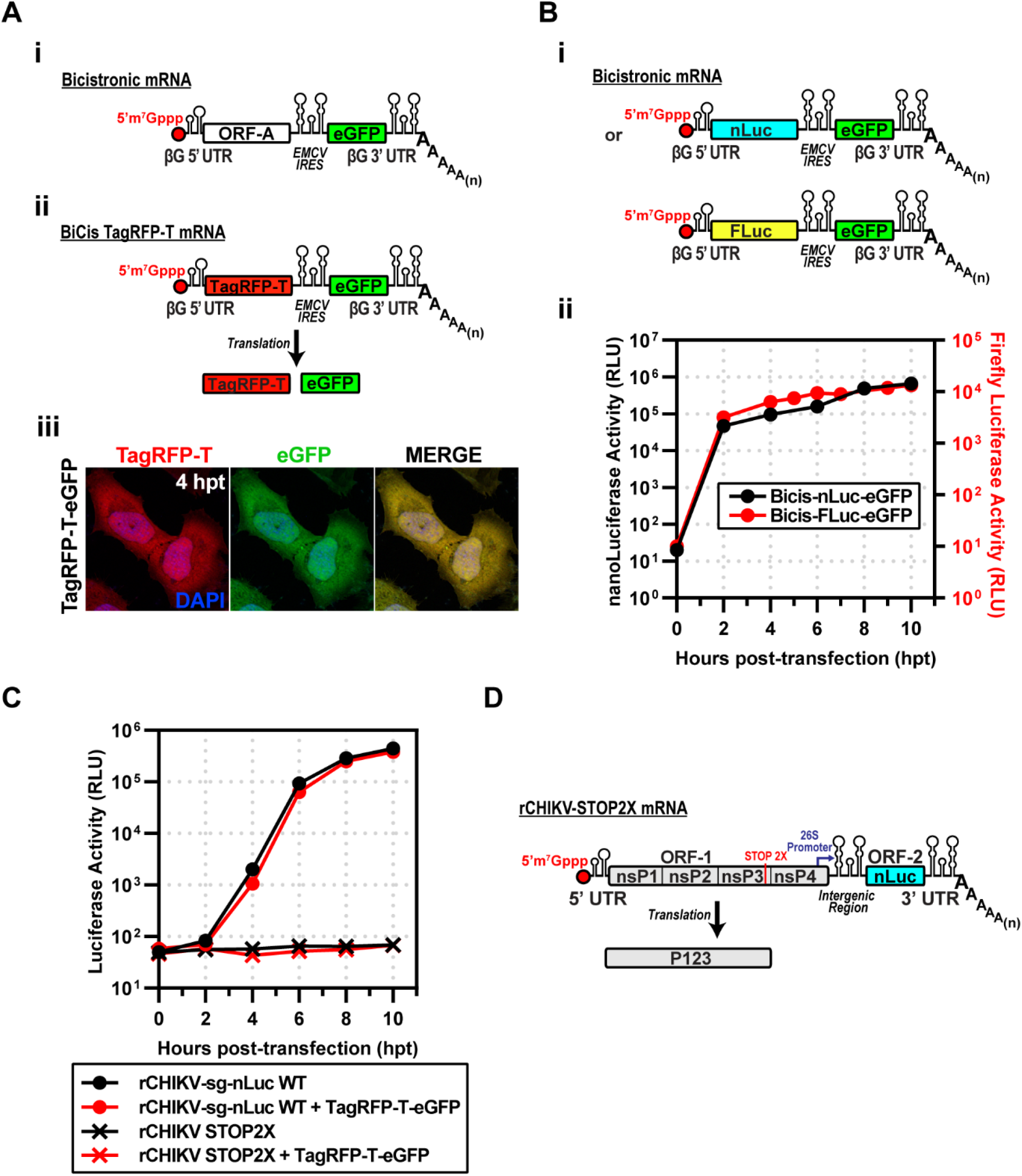
Strategy to express proteins in cells replicating subgenomic RNA replicons. **(A)(i)** Design of a bicistronic (BiCis) mRNA containing a β-globin 5′UTR, ORF-A, EMCV IRES, eGFP, β-globin 3′UTR, and poly(A). **(ii)** BiCis-TagRFP-T produces TagRFP-T (cap-dependent) and eGFP (IRES-dependent). **(iii)** Representative fluorescence images at 4 h post-transfection in HeLa cells. TagRFP-T (red) and eGFP (green) display a robust signal. **(B) (i)** BiCis nLuc and Fluc constructs, either reporter placed in ORF-A. **(ii)** Reporter translation confirmed by luciferase assays in electroporated HeLa lysates. **(C)** Replicon competition assays: nLuc activity of rCHIKV-sg-nLuc WT or STOP2X ± BiCis TagRFP-T/eGFP. WT replicons produced robust nLuc activity regardless of co-expression; STOP2X yielded no signal. **(D)** Schematic of rCHIKV STOP2X, with two stop codons replacing the opal site, preventing nsP4 expression, accordingly producing P123 only.

The second experiment we performed assessed the kinetics of gene expression after inserting genes into the first cistron. To do this quantitatively, we used luciferase reporters. To compare differences in gene sequences, we introduced either nLuc or firefly luciferase (FLuc) (panel i of **Fig. 2B**). Both reporters reached near maximum levels of expression by two hours post-transfection (hpt) (panel ii of **Fig. 2B**). The dynamic range of nLuc was 10-fold greater than observed for FLuc (panel ii of **Fig. 2B**).

The goal was to recapitulate the Lemm and Rice experiment, co-expressing P123 and P34. P34 would be expressed using the bicistronic mRNA. The possibility existed that co-transfection of the mRNA and the subgenomic replicon would compete for the translation machinery. To test this possibility directly, we co-transfected the CHIKV subgenomic replicon encoding nLuc with the BiCis TagRFP-T mRNA. The kinetics of luciferase activity were the same in the absence and presence of the BiCis mRNA (**Fig. 2C**). We conclude that there is no competition between these two RNAs when introduced into the cell.

Production of the P1234 precursor requires readthrough of an opal (UGA) stop codon at the 3’-end of the nsP3-coding sequence [4] . To construct a subgenomic mRNA that would only produce P123, we substituted the opal stop codon with ochre (UAA) and opal (UGA) stop codons. This mRNA is referred to as rCHIKV-STOP2X (**Fig. 2D**). Transfection of this mRNA into cells failed to produce any detectable luciferase activity (**Fig. 2C**).

### Expression of CHIKV P123 and P34 is sufficient for subgenomic RNA synthesis

We were now able to repeat with CHIKV the experiment performed by Lemm and Rice with SINV [7] . We co-transfected rCHIKV-STOP2X and BiCis-P34 mRNAs to produce P123 and P34 precursor proteins (**Fig. 3A**). These proteins supported minus-strand synthesis followed by genomic and subgenomic RNA synthesis (**Fig. 3B**). The subgenomic RNA encodes nLuc, so we followed subgenomic RNA synthesis indirectly by monitoring luciferase activity (**Fig. 3B**). Transfection of STOP2X mRNA alone yielded no luciferase signal (STOP2X in **Fig. 3C**). In combination with P34 mRNA, however, luciferase activity was observed beginning at 2 h post-transfection (STOP2X + P34 WT in **Fig. 3C**). The observed kinetics were on par with the kinetics observed for the complete CHIKV replicon (rCHIKV-sg-nLuc WT in **Fig. 3C**). Production of luciferase activity required the nsP4 domain of P34 to be an active RdRp as mutation of the signature GDD motif to GAA precluded production of luciferase activity (STOP2X + P34 GAA in **Fig. 3C**). We observed a dependency on the amount of P34 mRNA delivered with the STOP2X for complementation as assessed by luciferase activity (**Fig. 3D**). Inactivating nsP2-dependent proteolysis of P34 (**Fig. 3E**) eliminated complementation (**Fig. 3F**).

**Figure 3.**
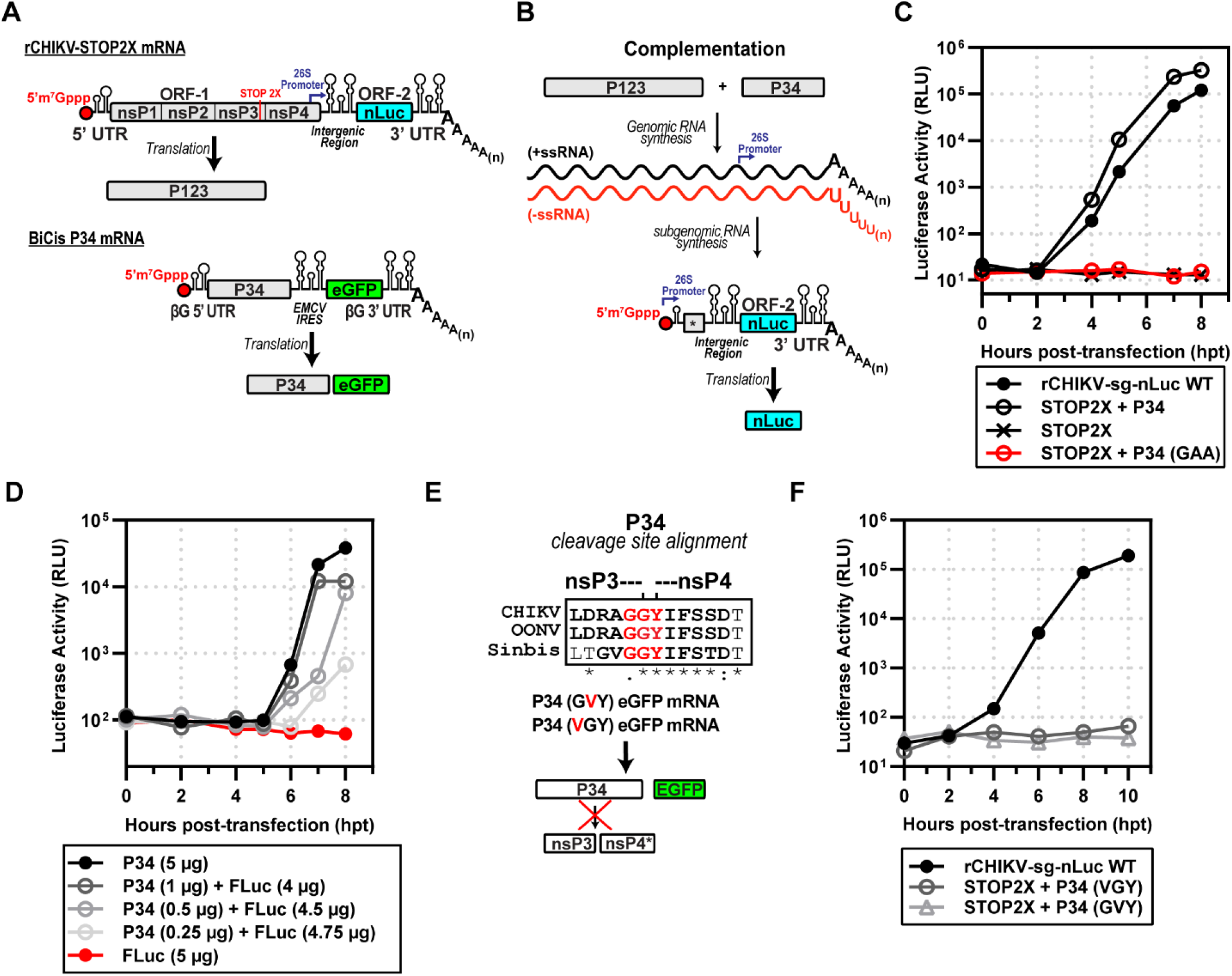
Expression of CHIKV P123 and P34 is sufficient for subgenomic RNA synthesis. **(A)** rCHIKV STOP2X lacks nsP4, producing P123 alone; BiCis P34 provides nsP4 in the form of P34 (plus eGFP marker). **(B)** Complementation assay schematic: co-transfection allows P123 + P34 to drive RNA replication and nLuc expression. **(C)** nLuc activity: STOP2X complemented with P34 WT restoring replication; P34-GAA (inactive RdRp) failed. **(D)** P34 titration: graded P34 RNA doses supported subgenomic RNA synthesis, with Fluc RNA included to equalize RNA input. **(E)** Mutagenesis of the nsP3-nsP4 conserved junction among alphaviruses prevents cleavage (VGY or GVY mutants). **(F)** These uncleavable P34 variants failed to complement STOP2X.

### Identification of a minimal P34 precursor for subgenomic RNA synthesis

The nsP3 domain of P34 is a multifunctional protein. At the amino terminus, there is a macro domain-containing mono-ADP-ribosylhydrolase activity (MD in **Fig. 4A**) [30], followed by an alphavirus unique domain of unknown function (AUD in **Fig. 4A**) [31], and ending with a hypervariable domain (HVD) that is an intrinsically disordered domain contributing to myriad interactions with the host that contribute positively to alphavirus multiplication (**Fig. 4A**) [32–34] . For SINV, the nsP3 domain of P34 is 549 amino acids [1] . Lemm and Rice showed that addition of only 22 – 153 amino acids from the carboxyl terminus of nsP3 to nsP4 was sufficient for complementation [7,12] . These truncated forms of P34 appeared to be equivalent functionally, at least within the sensitivity of the assays used [7,12] .

**Figure 4.**
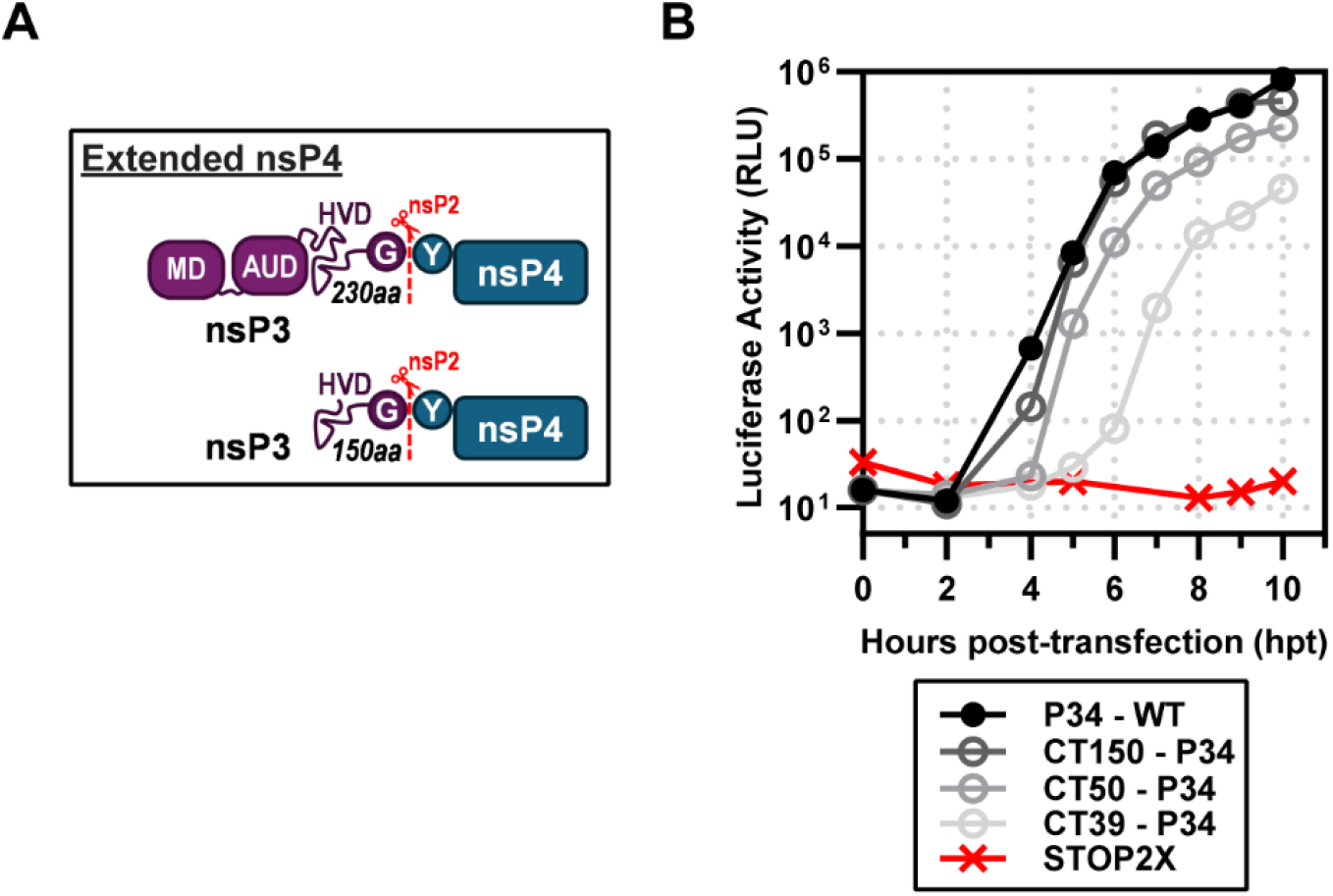
Identification of a minimal P34 precursor for subgenomic RNA synthesis. (**A)** Schematic of extended nsP4 constructs lacking domains of nsP3: macro domain (MD), alphavirus unique domain (AUD), and partial hyper-variable domain (HVD), produce CT150, CT50, or CT39-P34. **(B)** Complementation assays: STOP2X supported subgenomic RNA synthesis with P34 WT, CT150, and CT50, and partially with CT39-P34.

We performed a similar experiment for CHIKV. We started with a P34 derivative containing only the carboxy-terminal 150 amino acids of nsP3. This derivative was named CT150-P34. In addition to this derivative, we constructed CT50-P34 and CT39-P34. All of these derivatives exhibited some level of activity (**Fig. 4B**). Luciferase activity produced by CT150-P34 essentially superimposed with P34 (**Fig. 4B**). CT50-P34 exhibited a short lag in the accumulation of luciferase activity relative to P34 but reached very similar endpoints (**Fig. 4B**). CT39-P34 was clearly debilitated relative to P34, but it functioned. The CT50-P34 derivative is currently under investigation [18] .

### Ub-nsP4 fusions do substitute for P34 when expressed transiently

The early studies of Lemm and Rice showed that the authentic tyrosine amino terminus of nsP4 supports the most efficient synthesis of SINV RNAs [7,22] . Since that time, the lab of Andres Merits has reinforced this conclusion for many other alphaviruses [35] . To produce Tyr•nsP4, these investigators fused Tyr•nsP4 to ubiquitin (Ub) to create Ub-Tyr•nsP4 (**Fig. 5A**). Ubiquitin carboxy-terminal hydrolases remove Ub cleanly, likely co-translationally, to release Tyr•nsP4 (**Fig. 5A**) [36,37] . Unexpectedly, Ub-Tyr•nsP4 did not substitute well for P34 (**Fig. 5B**). Some luciferase activity was detected, but Ub-Tyr•nsP4 exhibited an extreme delay of at least 5 h before luciferase activity was detected (**Fig. 5B**). At 10 h post-transfection, this derivative was still at a level of luciferase activity more than 3-logs lower than observed for P34 (**Fig. 5B**). The activity for Ub-Tyr•nsP4 required ubiquitin cleavage as a cleavage-resistant derivative was dead (Ub*-Tyr•nsP4 in **Fig. 5B**). A pre-processed form of nsP4, Met•Tyr•nsP4, also failed to support RNA synthesis and consequentially luciferase activity (**Fig. 5B**). As discussed below, the inability of the Ub-Tyr•nsP4 to function in our system is likely attributable to the transient nature of expression and/or levels of the fusion that accumulate in cells.

**Figure 5.**
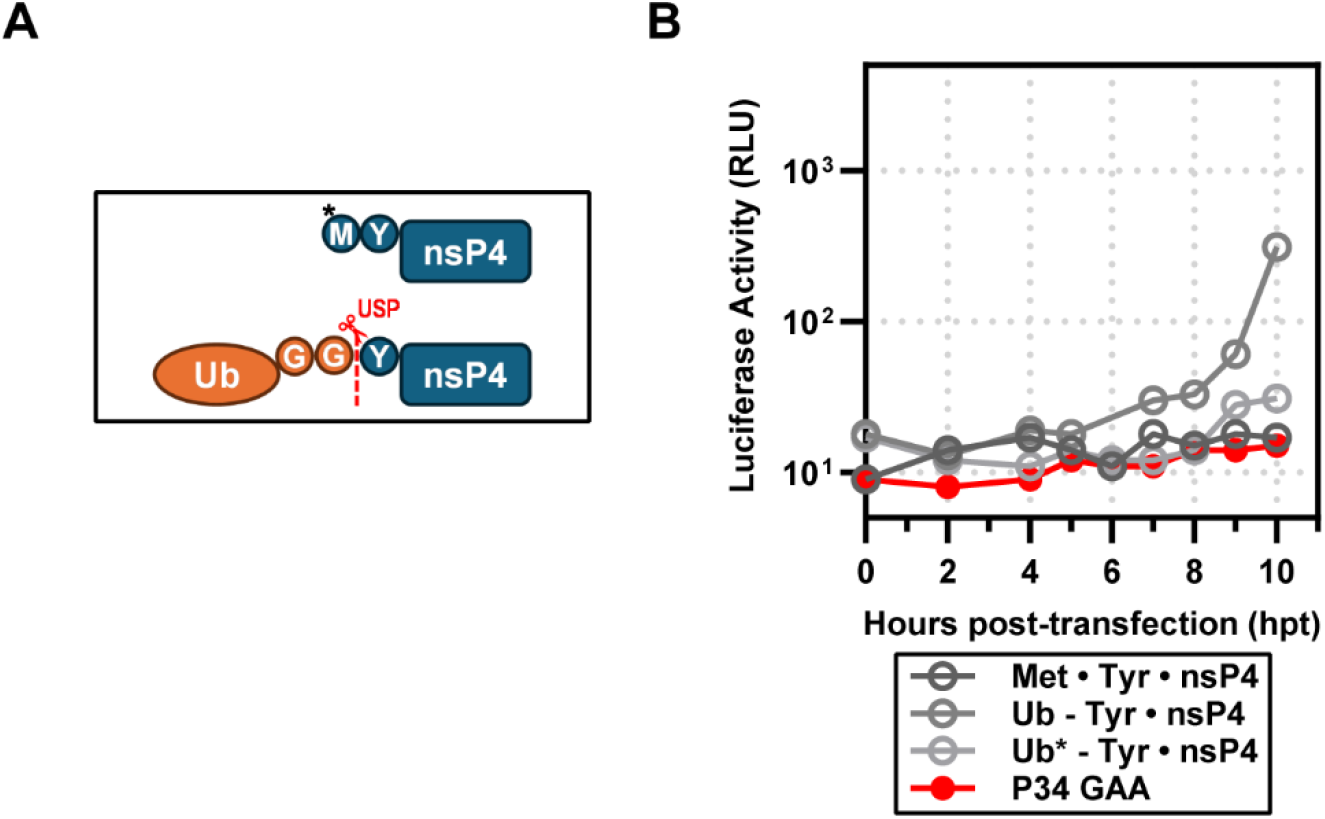
Ub-nsP4 fusions do substitute for P34 when expressed transiently. **(A)** Schematic of nsP4 constructs included: Met•Tyr•nsP4, Ub-Tyr•nsP4 (cleavable), and Ub*- Tyr•nsP4 (uncleavable). **(B)** Complementation assays: STOP2X was partially and inefficiently rescued by Ub-Tyr•nsP4 but not by Met•Tyr•nsP4 or Ub*-Tyr•nsP4.

## DISCUSSION

Alphaviruses are a persistent and growing threat to public health [38] . Epidemics of Chikungunya virus (CHIKV) are currently ongoing [39] . The safety of existing vaccines to prevent CHIKV infection and/or the sequelae associated with infection is questionable [40,41] . The development of antivirals to address infections caused by CHIKV and other alphaviruses is therapeutically needed. The nsP2-encoded protease [8,42], ATPase and helicase [43], and the nsP4-encoded RNA-dependent RNA polymerase (RdRp) [15,44] represent important targets for the development of antiviral therapeutics. The biochemical assays to study the nsP2-encoded activities are well established [10,45] . However, the ability to study kinetics, mechanism, and inhibition of the nsP4-encoded RdRp in vitro does not exist. The goal of this study was to identify a minimal form of CHIKV nsP4 that was active in a cell-based system, making it tractable for biochemical and biophysical characterization.

Early studies from the Strauss and Rice laboratories showed that a polyprotein precursor form of nsP4, specifically P34, an nsP3-nsP4 polyprotein, could support genome replication when co-expressed with the P123 polyprotein precursor [6,9] . Data from the Rice laboratory suggested that precursor forms of nsP4 were essential because of the need for an authentic tyrosine amino terminus of nsP4 for genome replication to occur [7,12,22] . Expression of a ubiquitin-nsP4 fusion complemented P123 as well as P34 in experiments from his lab [7,12] and others [16,17,46] .

However, nsP4 with an authentic amino terminus lacks any demonstrable RdRp activity in vitro [5,17,47] and, at best, has been shown to exhibit terminal transferase activity [5] . Biophysical studies performed by our laboratory using the O’nyong’nyong virus (ONNV) nsP4 RdRp with an authentic tyrosine amino terminus revealed an unexpected, extended conformation that explained the absence of demonstrable polymerase activity for this form of the protein [18] .

Previous studies evaluating the activity of nsP4 precursor proteins in genome replication have constitutively overexpressed these proteins either using a vaccinia virus expressing bacteriophage T7 RNA polymerase [7,13] or the cellular RNA polymerase II (RNAPII) directed by a cytomegalovirus (CMV) promoter [17,19,48] . The presence of vaccinia virus in a cell where alphavirus genome replication is being monitored was state-of-the-art in the 1990s but is now perceived as a confounding factor. The use of RNAPII requires the transfection of DNA and, minimally, tens of hours for expression to occur [19,49,50] .

We opted to use an RNA-based system. Delivery of viral or recombinant mRNAs to the cytoplasm results in expression within minutes, as observed during a bona fide viral infection. In addition, the technology to suppress, if not eliminate, activation of innate immune sensors by transfected mRNA continues to improve [49,51–53] .

Transfection of CHIKV mRNA with an authentic, capped, AU 5’-end encoding P1234 and nLuc in place of the coding sequence for the structural proteins yielded a near 5-log increase in luciferase over a 10 h period (**Fig. 1**). We constructed a bicistronic mRNA to use for expression of nsP4 derivatives that we validated by inserting coding sequence for two different luciferase enzymes (**Fig. 2**). Co-expression of the CHIKV P123 precursor instead of the P1234 precursor and the CHIKV P34 precursor was sufficient to produce levels of luciferase activity equivalent to the primary CHIKV replicon (**Fig. 3**). Luciferase activity correlated linearly to the amount of P34 expressed to a limit (**Fig. 3**). Cleavage of P34 by the nsP2-encoded protease was essential to observe luciferase activity (**Fig. 3**). Removal of all nsP3 except for the 50 carboxy-terminal residues (CT50-P34) supported near wild-type levels of luciferase activity (**Fig. 4**).

Expression of P34 in bacteria results in substantial cleavage of the protein by bacterial proteases, which target the very large, intrinsically disordered domain (our unpublished observations). Once we learned that CHIKV CT50-P34 functioned in cells, we expressed and purified ONNV CT50-P34 [18] . Unlike nsP4, which forms an extended conformation, CT50-P34 forms a compact conformation and assembles into tetramers [18] . Our working hypothesis is that P34 and functional derivatives thereof bind to the central channel of an nsP1 dodecamer (**Fig. 6A**). nsP2 cleaves the (nsP1)12-bound P34 activating its RdRp activity (**Fig. 6B**). Physical interactions between nsP4 and (nsP1)12 maintain nsP4 in its active conformation (**Fig. 6B**). Once RNA synthesis is complete, dissociation of nsP4 from (nsP1)12 causes a fold switch to the extended, inactive conformation (**Fig. 6C**). This change may prevent the RdRp from using cellular RNA to produce dsRNA that would activate intrinsic defense mechanisms and innate immune responses of the cell [54–56] .

**Figure 6.**
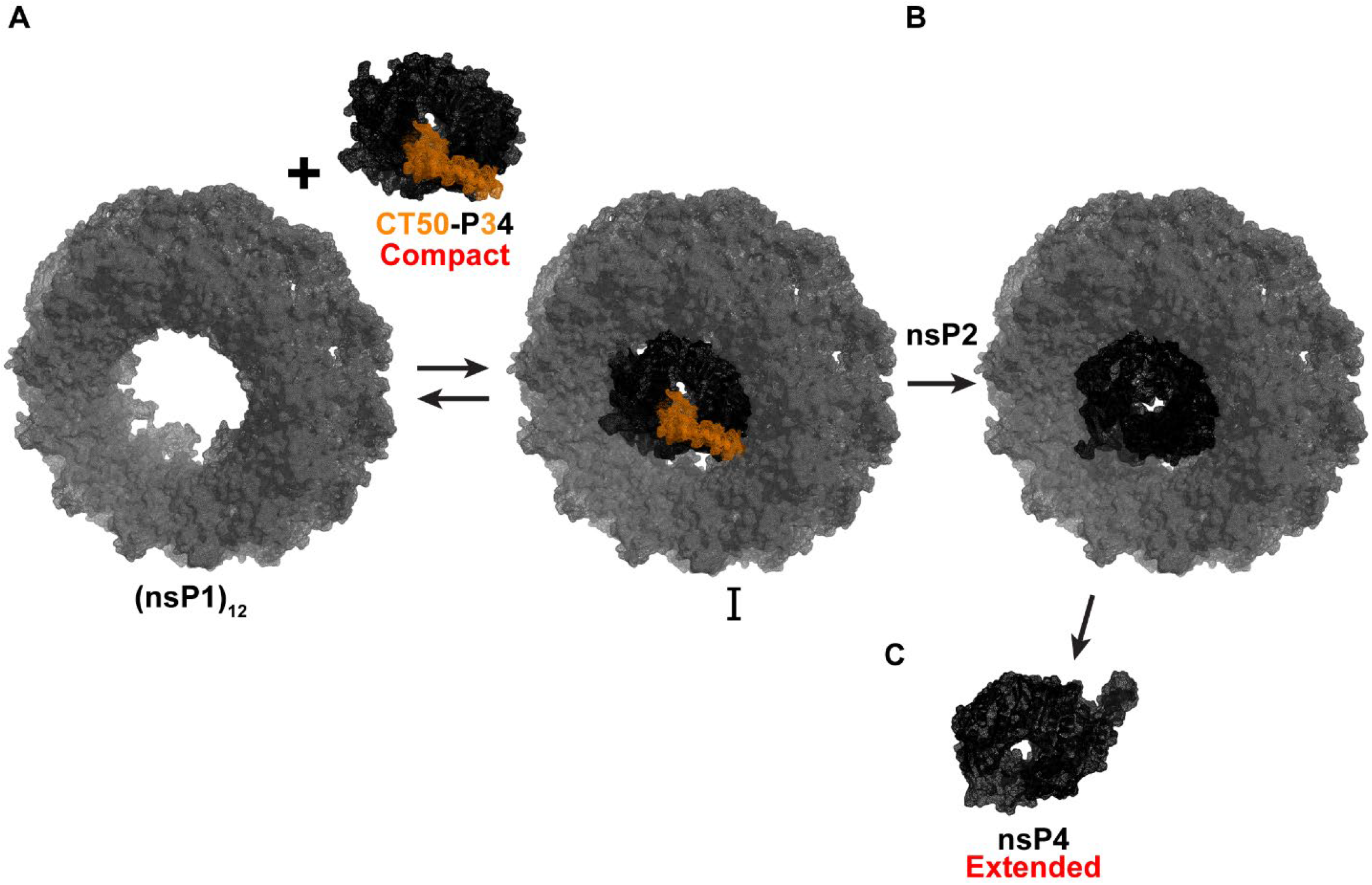
Proposed model for an nsP4 precursor role in CHIKV RNA synthesis. **(A)** P34 and functional derivatives (CT50-P34) bind to the central channel of an nsP1 dodecamer. (B)nsP2 cleaves the (nsP1)_12_-bound P34, activating its RdRp activity. Physical interactions between nsP4 and (nsP1)_12_ maintain nsP4 in its active compact conformation. **(C)** Once RNA synthesis is complete, dissociation of nsP4 from (nsP1)_12_ causes a fold switch to the extended, inactive extended conformation.

The one unexpected observation of the experiments reported here was the inability to have a Ub-nsP4 fusion substitute for P34, as observed by other laboratories (**Fig. 5**). Ub fusions are generally co-translationally processed unless sufficient fusion protein is made to overwhelm the ubiquitin-specific carboxy-terminal hydrolases [7,17,37] . Expression of Ub-nsP4 using the bicistronic mRNA approach may be high initially but will diminish over time because the mRNA will be degraded, and nsP4 will undergo turnover [55]. The systems in which Ub-nsP4 has been shown to work constitutively express the fusion protein, which is the simplest explanation for the difference between the systems.

In conclusion, we have established an RNA-based replicon and trans-complementation system for CHIKV that should apply to alphaviruses broadly. The primary benefit of this approach is that it is faster than DNA-based approaches. Using this system, we have defined the CT50-P34 precursor as the minimal donor of CHIKV nsP4 RdRp activity, consistent with findings for SINV [7,18,57] . The capacity to purify the ONNV CT50-P34 derivative [18] bodes well for the purification of CHIKV CT50-P34. The ability to utilize biological, biochemical, and biophysical approaches to study CHIKV genome replication represents a significant advancement that will likely fill gaps in our understanding of alphavirus genome replication that have existed for decades.

## LIMITATIONS OF THE STUDY

We have been unable to directly quantify the expression of nsP4 or polyproteins thereof because all antibodies against nsP4 have exhibited very high backgrounds. This circumstance is likely to change with the use of our purified preparations of nsP4 and CT50-P34 as antigens. We can also try to add immunological tags to the nsP4-coding sequence.

We are only monitoring subgenomic RNA synthesis using the replicons reported in this study. The assumption is that minus- and full-length genomic RNA synthesis are also occurring. While minus-strand synthesis cannot be monitored using reporters, future versions of the reported replicon will benefit from addition of a reporter to the P123 open reading frame.

## PERSPECTIVES

The RNA-based replicon used here will permit a rapid assessment of the determinants of the CHIKV genome required for RNA synthesis. The system reported herein should enable the discovery of determinants of CT50-P34 required for complementation. Additionally, interactions between various forms of nsP4 and other non-structural proteins can be examined. Addressing these fundamental questions may permit reconstitution of the CHIKV replisome in vitro from purified components and pave the way for the first mechanistic description of nucleotide addition by an alphavirus polymerase.

## MATERIALS AND METHODS

### Cell culture

HeLa cells (CRM-CCL-2; ATCC) were maintained in Dulbecco’s Modified Eagle Medium: Ham’s F-12 nutrient mixture (DMEM/F12; Gibco) supplemented with 10% heat-inactivated fetal bovine serum (HI-FBS; Atlanta Biologics) and 1% penicillin-streptomycin (P/S; Corning). Cells were cultured at 37 °C in a humidified incubator with 5% CO_2_.

### Plasmids

Insertions and deletions were generated by overlap extension PCR or synthesized as gBlocks (Integrated DNA Technologies, IDT). Constructs were assembled using In-Fusion seamless cloning (Takara). All insertions, deletions, or point mutations were verified by Sanger sequencing (short fragments) or Oxford Nanopore long-read sequencing (whole plasmids). rCHIKV constructs derived from published sequences (GenBank accessions GU189061 and EF452493). Bicistronic constructs incorporated the human β-globin sequence (U01317) and the EMCV IRES, as described by Bochkov et al. (2006) [28] .

### In vitro transcription

Plasmids encoding rCHIKV genomes (full-length or replicon) were linearized with NotI, and BiCis constructs with SalI positioned immediately downstream of the encoded poly(A) tail. Linearized templates were transcribed in vitro using T7 RNA polymerase produced in-house (Cameron Lab). Reactions were treated with 2 U Turbo DNase (ThermoFisher) to remove residual DNA. rCHIKV RNAs were co-transcriptionally capped with CleanCap AU (TriLink), while BiCis RNAs were capped with CleanCap AG (TriLink). Transcripts were purified with the RNeasy Mini Kit (Qiagen), quantified spectrophotometrically, and resuspended in RNase-free water. Purified RNAs were delivered into cells by electroporation (Gene Pulser, Bio-Rad).

### Subgenomic replicon luciferase assay

Luciferase assays were performed as described previously [58] . Briefly, 5 µg of in vitro– transcribed replicon RNA was electroporated into HeLa cells, which were then cultured in standard growth medium. At the indicated times post-electroporation, cells were harvested and lysed in 100 µL of 1× Cell Culture Lysis Reagent (CCLR; Promega).

Luciferase activity, as a surrogate for subgenomic RNA replication, was measured in lysates after addition of an equal volume of Nano-Glo or Firefly Luciferase substrate (Promega). Luminescence was quantified using either a Junior LB 9509 luminometer (Berthold Technologies) or a BioTek plate reader.

### Antibodies

Primary antibody: rabbit anti-α-tubulin (Cell Signaling; 1:500). Secondary antibody: goat anti-rabbit IgG (H+L) conjugated to Alexa Fluor dyes (Invitrogen; 1:1000), used for immunofluorescence.

### Immunofluorescence

Immunofluorescence assays were performed as described previously [59] . HeLa cells were grown on glass coverslips, treated as described in the respective figure legends, and fixed at the indicated time points with 4% paraformaldehyde in PBS for 20 min. For UnaG reporter detection, cells were supplemented with 7 µM bilirubin (APEX Bio) to enhance fluorescence. Cells were permeabilized with 0.2% Triton X-100 in PBS (10 min), blocked with 3% goat serum (1 h), and incubated with primary antibodies (1 h). Following washes, coverslips were incubated with secondary antibodies (1 h) and counterstained with either DAPI (Sigma) or TO-PRO-3 (Invitrogen) for 10 min. Coverslips were mounted in ProLong Glass Antifade Mountant (ThermoFisher).

### Imaging

Fluorescence imaging was performed on a BZ-X800 all-in-one microscope (Keyence) using a 40× objective. Images were processed with BZ-X800 Analyzer software (Keyence). Multiple images were acquired, and representative cells were selected for display.

## AUTHOR CONTRIBUTIONS

Conceptualization and/or Design of Research (DAD, JJA, CEC); Data Collection and Acquisition (DAD, AJ, JMH); Data Analysis (DAD, AJ, CEC): Data Interpretation: (DAD, JJA, CEC); Contributed New Reagents or Analytical Tools: (DAD, AJ, JMH); Visualization & Figures (DAD, CEC); Writing original draft (DAD, CEC); Writing – review & editing (DAD, AJ, JMH, JJA, CEC); Supervision (DAD, CEC); Project Administration (DAD, CEC); Funding Acquisition (JJA, CEC).

## ACKNOWLEDGMENTS

The corresponding authors thank members of our lab and collaborators for their feedback in the progression of these studies. The authors also thank Che-Kang Chang and Mark Heise for sharing plasmids and the original sequences of CHIKV.

## SUPPLEMENTARY DATA

Supplementary data can be made available upon request.

